# A point mutation in the nuclear speckle protein and splicing factor SRRM2 is associated with amyotrophic lateral sclerosis and causes dysregulation of synapse-associated genes

**DOI:** 10.1101/2025.09.11.675713

**Authors:** Qingyu Shi, Chloe Lauder, Jolie M. Miller, Yong-Dong Wang, Noha Elsakrmy, Sofija Volkanoska, Brian D. Freibaum, Hui Peng, Hong Joo Kim, J. Paul Taylor, Haissi Cui

**Affiliations:** Department of Chemistry, University of Toronto, Toronto, Ontario M5S 3H6, Canada; Department of Cell and Molecular Biology, St. Jude Children’s Research Hospital, Memphis, TN, USA; School of the Environment, University of Toronto, Toronto, Ontario, Canada; Structural Genomics Consortium, University of Toronto, Toronto, Ontario, Canada

## Abstract

Amyotrophic lateral sclerosis (ALS) is a fatal neurodegenerative disorder characterized by the progressive loss of motor neuron function. ALS is a multifactorial disease which can originate from complex genetic and environmental factors. The identification of risk factors and their molecular contribution to ALS expand our understanding of the disease mechanism.

Here, we describe a family with dominantly inherited degeneration, which carries a mutation in the serine/arginine repetitive matrix 2 gene (*SRRM2)*. SRRM2 is essential for nuclear speckle formation and a constitutive member of the RNA splicing machinery. To investigate how the mutation in SRRM2 contributed to the ALS pathogenesis, we examined its effect on a model cell line, where the point mutation was introduced in the endogenous gene. Surprisingly, we found that the resulting single amino acid exchange led to the loss of one protein-protein interaction, between SRRM2 and the splicing factor ACIN1. Transcriptome studies further revealed wide-spread differential gene expression, which converged on the dysregulation of synapse-associated pathways. Together, our findings identify SRRM2 as a novel ALS risk factor and provide mechanistic insights into how its mutation can be linked to ALS pathology.

## Introduction

Neurodegenerative disorders are characterized by the progressive loss of nerve structures and synaptic connections, which can affect the central and peripheral nervous system of the brain’s function (1, 2). They arise from a wide range of biological and environmental factors, including genetics, cell signaling impairments, neuronal apoptosis, inflammatory responses, and aging (1, 3). Prominent neurodegenerative disorders include Alzheimer’s disease, Parkinson’s disease, Huntington’s disease, multiple sclerosis, and amyotrophic lateral sclerosis (ALS) (1, 4). ALS expresses rapidly fatal neurological conditions, characterized by degeneration of upper and lower motor neurons in both the brain and the spinal cord. This disease represents a heterogenous group of both familial (∼10%) and sporadic (∼90%) forms (3). To date, there are over 100 candidate genes that have been reported, with 16 of them identified as causative for ALS pathogenesis (3, 5). These genes largely fall into three functional categories: genes that regulate proteostasis and protein quality control, factors that are involved in RNA processing and assembly into membrane-less granules, and components in the cytoskeleton or other facilitators of intracellular trafficking (3). Importantly, dysregulation of nuclear speckles in ALS can reduce global mRNA splicing efficiency and broadly affect RNA splicing in the ALS patients (6).

Nuclear speckles are membrane-less organelles which are located in the interchromatin space of the nucleus (7). They are dynamic structures that appear as irregularly shaped clusters of varying sizes under microscopy. Their molecular constituents exchange rapidly with the surrounding nucleoplasm (7, 8). One of the core components of nuclear speckles is Serine/Arginine Repetitive Matrix Protein 2 (SRRM2), a nuclear SR protein essential for pre-mRNA splicing and nuclear speckle formation (9, 10). SRRM2 is a spliceosomal component that enhances splicing efficiency via facilitating interactions between pre-mRNA and the spliceosomal catalytic machinery (10, 11). It also interacts with other SR-related proteins, most notably SON, a protein splicing cofactor (11). Importantly, co-depletion of both SRRM2 and SON results in a near-total breakdown of nuclear speckles, highlighting that both proteins are essential nuclear speckle scaffolds (6, 12). Structural studies suggest that the conserved N-terminal RNA recognition motif of SRRM2 directly binds mRNA, while the intrinsically disordered regions (IDRs) promote multivalent protein-protein interactions (12, 13). These IDRs are enriched in serine/arginine residues, which drive liquid-liquid phase separation, a biophysical process that is essential for speckle assembly (13).

Recent studies showed that the dynamic three-dimensional organization of genomic DNA around nuclear speckles is crucial for efficient mRNA processing (14). Further, nuclear speckles act as concentrated reservoirs for splicing factors to engage nascent pre-mRNAs (9, 15, 16). The spatial proximity of transcription relative to the speckle can influence mRNA transcription and processing efficiency: pre-mRNA located near speckles encounters a higher concentration of splicing factors, while in contrast, pre-mRNAs located farther from the speckles are exposed to a lower concentration of splicing factors, resulting in less efficient splicing (14). Nuclear speckles can thereby regulate splicing by promoting spatial proximity and increasing the concentration of mRNA processing factors relative to RNA (17–19). In addition to RNA processing, RNA transcription is increased in the proximity of nuclear speckles (20). Further, nuclear speckles store mRNA transcripts upon transcription inhibition to later release them (21). While in the nuclear speckle, these transcripts are protected from degradation (21), suggesting a regulatory function of nuclear speckles in mRNA transcript abundance, availability, and lifetime. Altogether, these mechanisms establish nuclear speckles as active regulatory components that play a decisive role in RNA transcription and processing.

Recently, nuclear speckles, and by extension SRRM2, have been implicated in ALS disease progression. Nuclear speckle integrity is disrupted by repeat expansions in C9ORF72, one of the best studied genetic drivers of ALS (6). The (GGGGCC)n repeat RNA has been shown to co-localize with nuclear speckles and alters their dynamics and phase separation behavior (6, 22, 23). Additionally, one of the protein products of C9ORF72-repeat expansions, poly-GR stretches, interact with SRRM2, and poly-GR aggregates can sequester SRRM2 into cytoplasmic inclusions (6, 24, 25). Together, these processes contribute to nuclear speckle dysfunction and result in extensive alternative splicing abnormalities, including global exon skipping and intron retention, ultimately leading to neuronal degeneration (6).

Genetic variants of SRRM2 have been reported in other neurological diseases, including Parkinson’s disease, papillary thyroid carcinoma, and neurodevelopmental disorders (26–28), suggesting that its role in RNA regulation is critical. Importantly, RNA processing defects are one of the key drivers of ALS, and mutations in various RNA-binding proteins have been linked to ALS disease pathology. Mutations in heterogeneous nuclear ribonucleoprotein A1 (hnRNPA1) lead to aberrant neuronal RNA splicing and result in neurodegeneration (29). Mutations in TAR DNA binding protein 43 (TDP43) cause cytoplasmic aggregation, while mutations in Fused in Sarcoma (FUS) result in widespread RNA splicing defects, changes in synaptic function, and FUS aggregates (30, 31). Overall, these genetic and functional associations demonstrate the importance of RNA processing in ALS disease progression.

Here, we describe a mutation in SRRM2, which was found in a family with ALS, 4089T.C, resulting in the exchange of a serine for a proline in position 1444 (S1444P). We established a cell line model of this variant to study SRRM2 S1444P and found that while nuclear speckle integrity was preserved, SRRM2 showed subtle but significant differences in its co-localization with SON. Interactome studies revealed the loss of a single interactor, the splicing factor ACIN1, which was confirmed by immunofluorescence. RNA-seq analysis suggested gene expression changes leading to the dysregulation of synapse-associated pathways. We therefore propose that SRRM2 mutations as a risk factor for ALS, with wide-spread consequences for cellular gene expression.

## Results

### Identification of an ALS-associated mutation in the nuclear speckle protein and splicing factor SRRM2

We identified a family with dominantly inherited degeneration characterized by cognitive impairment, motor neuron disease, Paget’s disease of the bone, and unspecified weakness, in whom mutations in known MSP-associated genes were absent (Figure 1A). Whole-exome sequencing was performed on samples from three affected individuals and one unaffected family member spanning two generations. After applying stringent filtering criteria and confirming segregation by Sanger sequencing in two additional family members, we identified five novel single-nucleotide variants in four genes that co-segregated with disease in this family (Supplementary Figure 1): *SRRM2*, WNT1 inducible signaling pathway-2 (*WISP2*), exostosin-2 (*EXT2*), and two variants in probable tubulin polyglutamylase TTLL2 (*TTLL2*).

**Figure 1.**
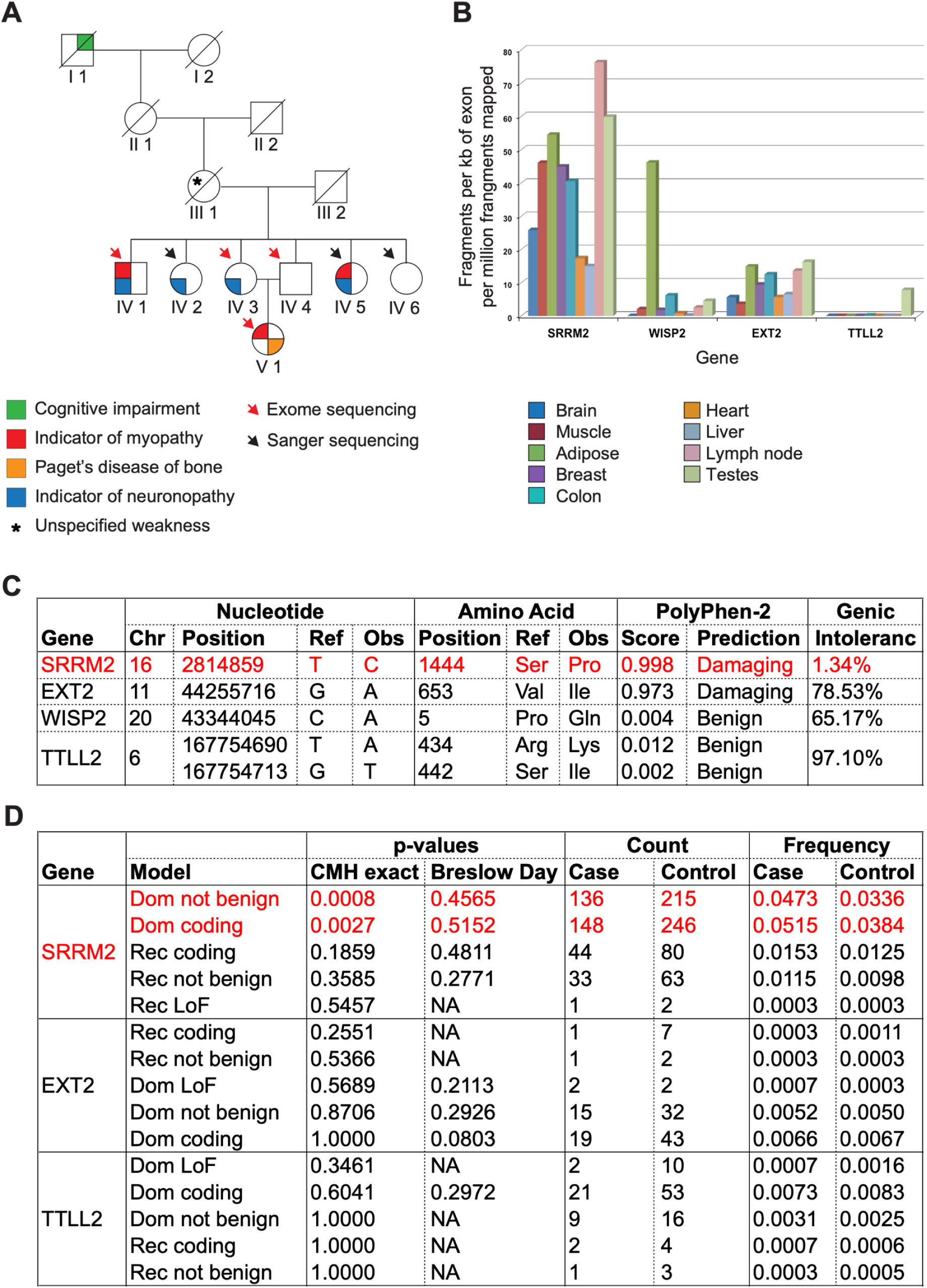
Identification of SRRM2 mutations as a potential ALS-risk factor. (A) Pedigree of an MSP family affected by ALS. Roman numerals indicate generations; Arabic numerals denote individuals within each generation. (B) The relative expression levels of SRRM2, WISP2, EXT2, and TTLL2 in multiple tissues are indicated as determined by RNA-seq. (C) Summary table of data relating to the 5 SNVs that co-segregate with disease in family. (D) Comparison of variant frequencies in the identified genes among 6,405 controls and 2,869 ALS patients from a published large-scale exome sequencing study (33). (A-D) MSP: Multisystem proteinopathy, SNVs: single-nucleotide variants

To prioritize candidate genes, we examined their expression patterns in disease-relevant tissues. SRRM2 was highly expressed in brain and muscle, both tissues relevant to MSP, while EXT2 showed moderate expression, and WISP2 and TTLL2 were not appreciably expressed in these tissues (Figure 1B). In silico analysis using PolyPhen-2 (32) predicted that the variants in *SRRM2* and *EXT2* were likely to be damaging (Figure 1C). Furthermore, evaluation of genic intolerance revealed that SRRM2 is highly intolerant to genetic variation (1.34%) compared to *EXT2* (78.53%) (Figure 1C), further implicating *SRRM2* as the most probable disease-causing gene in this family.

Notably, a large-scale exome-sequencing study comparing the frequency of variation in 17,249 genes between 6,405 controls and 2,869 patients also identified *SRRM2* as an ALS risk factor gene (33) (Figure 1D). Based on these findings, we focused subsequent investigations on exploring the potential disease connection between ALS and the SRRM2 variant, S1444P.

### Establishing a model cell line expressing the SRRM2 S1444P variant

In order to understand the molecular underlying as well as to confirm the pathogenicity of the S1444P variant of SRRM2, we decided to construct a model cell line to study alterations in SRRM2 interactome, nuclear organization, and mRNA splicing patterns. As overexpression of disordered proteins can lead to their mislocalization, we opted for genetic edits. We previously engineered a HEK 293T cell line expressing endogenously mVenus-labeled SRRM2 (19), where the yellow fluorescent protein mVenus is fused to the C-terminus of SRRM2, skipping the last 4 amino acids. This allows us to monitor and enrich endogenous SRRM2 through a fluorescent protein tag. As this cell line has already been extensively characterized (19), we opted to use this system as a starting point to investigate disease-associated variants of SRRM2. While HEK 293T cells are an imperfect model for a predominantly neurological disorder, they do show neuronal traits (34) and allow the facile investigation of nuclear and subcellular organization.

We chose prime editing to introduce the disease-causing 4089T.C SRRM2 mutation (35). While in the patient family, the variant was found as a de novo mutation in one allele only, we opted for a homozygous cell line to enable molecular studies of SRRM2 S1444P, which would allow us to attribute any observed changes to the mutation. To this end, we introduced gRNAs and repair templates to the Prime editing all in one (PEA1) (36) plasmid, which was modified with a mCherry fluorescent protein marker (Figure 2A). In order to increase efficiency, we co-transfected cells with pCMV_PE6c, an optimized prime editor (37). We then sorted for mVenus+/mCherry+ cells, which contained both the mVenus label on the SRRM2 C-terminus and were successfully transfected with PEA1_mCherry (Figure 2B). We recovered 15 colonies of which two were heterozygous and three were homozygous for the *SRRM2* 4089T>C mutation, as found by Sanger sequencing (Figure 2C). We confirmed these results with nanopore sequencing following PCR amplification (Supplementary Figure 2A) and continued our study with two of the homozygous cell lines. Cell proliferation was unchanged between wt and SRRM2 S1444P cells (Supplementary Figure 2B). SRRM2 mRNA levels were not robustly reduced but showed a trend to lower levels in homozygous S1444P cells (Figure 2D, Supplementary Figure 2C). On protein level on the other hand, SRRM2 was significantly decreased in S1444P homozygous cells, as shown by cytometry-based quantification of the C-terminally fused mVenus reporter protein (Figure 2E, gating scheme refers to Supplementary Figure 2D).

**Figure 2.**
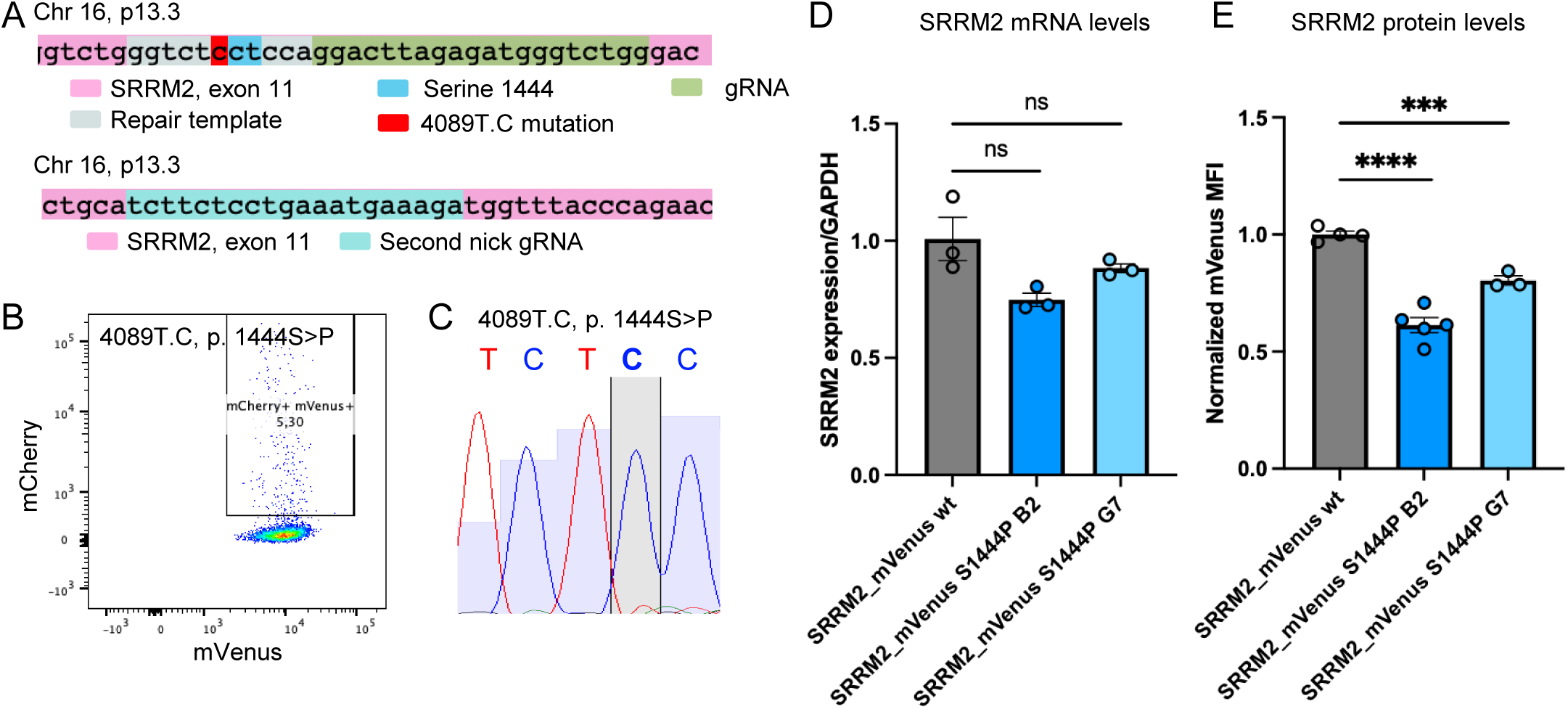
Establishment and characterization of a SRRM2 model cell line. (A) Scheme of gene editing design to exhange c.4089.T to C (S1444P) in the endogenous gene locus of SRRM2. (B) Verification of transfection and single cell sorting of HEK293T cells that were positive for both mVenus (C-terminal of SRRM2) and mCherry, denoting successful transfection using FACS. (C) Confirmation of a homozygous mutation in position c.4089.T>C through Sanger sequencing. (D) qPCR to assess SRRM2 mRNA levels following gene edit. (E) Flow cytometry to quantify SRRM2 protein levels based on SRRM2_mVenus fluorescence. (A-E) S1444P: amino acid exchange in SRRM2. wt: wildtype. (D, E) Mean with s.e.m, biological replicates, n=3, 3, 3, n=4, 5, 3. Unpaired, two-tailed t-test. ns=p>0.05. ***p<0.0002, ****p<0.0001.

### S1444P abrogates the interaction between SRRM2 and the splicing and apoptosis factor ACIN1

In order to investigate whether S1444P influences SRRM2’s interactome, we used a nanobody-mediated pulldown of SRRM2 under mild conditions to preserve interactions and assessed the resulting interactome using mass spectrometry. To this end, GFP-binding nanobody, which also recognized mVenus, was expressed and purified in *E. coli* and coupled to cyanogen bromide-activated magnetic Sepharose beads (Supplementary Figure 3A) (38). Cell lysates were then incubated with these beads overnight and proteins were eluted by trypsin digest. Cells expressing mVenus only were used as a control. Label-free quantification was performed to identify enriched proteins (Figure 3A, Supplementary Table S1, S2). When assessing the SRRM2 interactome (Figure 3B, left panel, Supplementary Table S1, SRRM2_mVenus vs mVenus), the largest enrichment was observed for SRRM2 and known interaction partners such as SON and SRRM1 were successfully retrieved (Figure 3B), confirming our approach. As expected, the majority of proteins found in the SRRM2 interactome were nuclear speckle proteins and mRNA splicing factors (Supplementary Figure 3B-E). We identified core spliceosomal proteins, including U2AF1, which recruits the U2 snRNP to branch point during splicing, ATP-dependent RNA helicases, such as DHX8, and exon-junction complex-associated proteins, including Pinin and EIF4A3.

**Figure 3.**
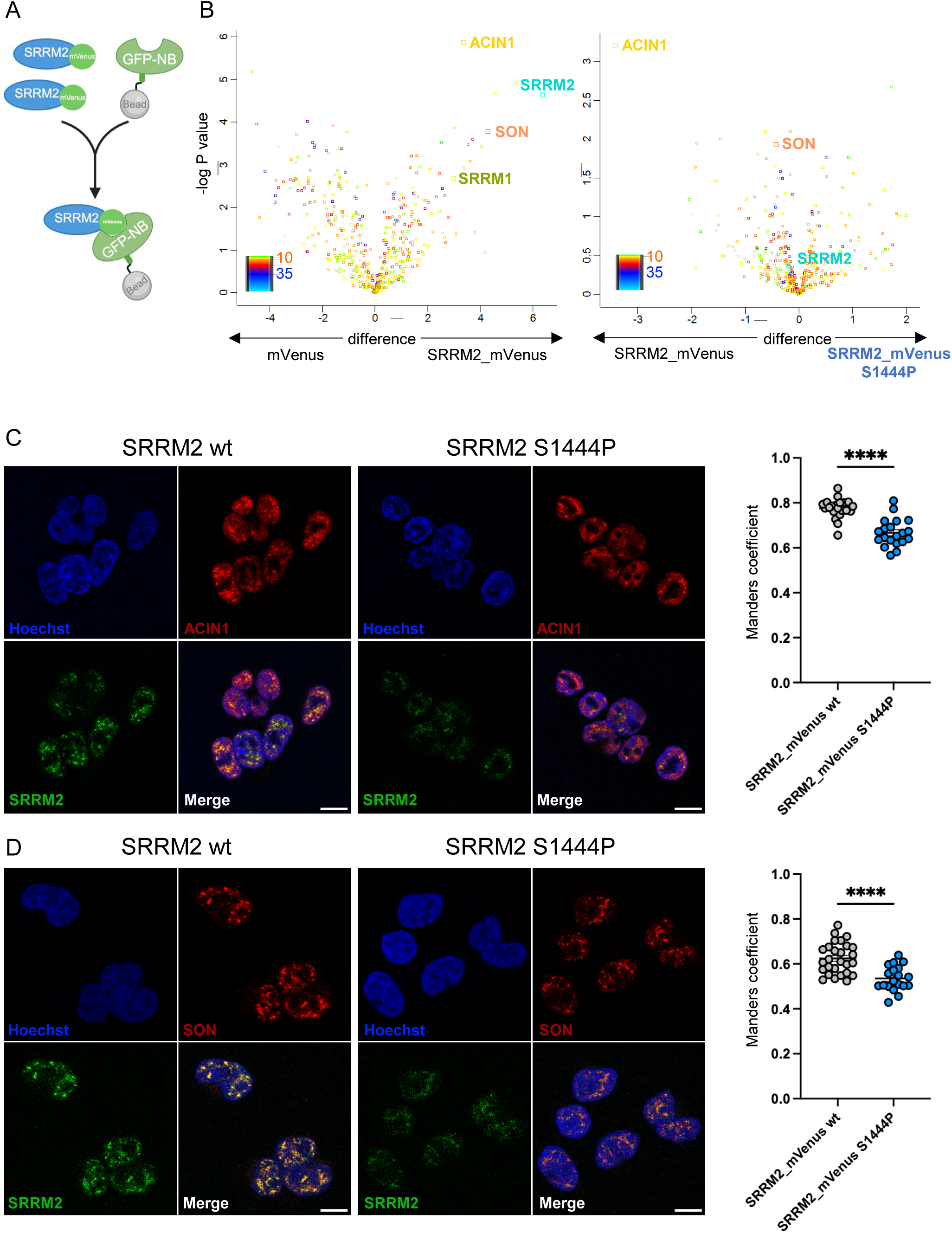
S1444P changes the SRRM2 interactome. (A) Scheme depicting SRRM2-mVenus pull-down with a GFP nanobody coupled to magnetic beads. mVenus: mVenus fluorescent protein with sequence similarity to GFP. GFP-NB: GFP nanobody. B) Left panel: Enrichment of SRRM2 and its interaction partners in SRRM2-mVenus cells following co-IP and mass spectrometry. Right panel: Comparison of the interactome between SRRM2 wt and S1444P cells. ACIN1 is highlighted. C) Left panel: Immunofluorescence of ACIN1 and SRRM2 in SRRM2 wt and S1444P cells, Right panel: Manders coefficient of SRRM2 colocalization with ACIN1. D) Left panel: Immunofluorescence of SON and SRRM2 in SRRM2 wt and S1444P cells, Right panel: Manders coefficient of SRRM2 colocalization with SON. (A-E) S1444P: amino acid exchange in SRRM2. B) Mass spectrometry was performed in biological triplicates. (C) Unpaired, two-tailed t-test: ****p<0.0001. A representative image of at least 2 replicates is shown. n= 21, 20. (D) Unpaired, two-tailed t-test: ****p<0.0001. A representative image of at least 3 replicates is shown. n= 27, 19

When comparing wt SRRM2 with the S1444P SRRM2 interactome, the most striking difference was the loss of Apoptotic chromatin condensation inducer in the nucleus (ACIN1) from the S1444P interactome (Figure 3B, right panel, Supplementary Table S2). ACIN1 is a heavily spliced gene, which is predominantly translated to a 152 kDa protein, and plays a role in mRNA splicing and apoptosis regulation (39, 40). Specifically, ACIN1 has been identified as a part of the exon-junction complex and it regulates DNA fragmentation by controlling splice isoform usage of key regulators (39, 41). Immunofluorescence analysis placed both SRRM2 and ACIN1 in the cell nucleus (Figure 3C, additional images Supplementary Figure 4A). We used colocalization to assess the degree of interaction. SRRM2-ACIN1 colocalization was decreased in S1444P cells as indicated by a lower Manders’ coefficient and vice versa (Figure 3C, right panel, Supplementary Figure 4B). While overall SRRM2 patterns were comparable between wt and S1444P, we were interested in probing whether more subtle changes occurred.

We therefore assessed the degree of colocalization between SRRM2 and SON, which are both nuclear speckle scaffolds (Figure 3D, additional images in Supplementary Figure 4C). In S1444P cells, colocalization of SRRM2 with SON decreased by 15% (Figure 3C) and vice versa (Supplementary Figure 4D). This suggests that the decrease in SRRM2 was reflected in the reduced colocalization with SON and/or subtle changes in nuclear speckle architecture were found in cells expressing SRRM2 S1444P.

As ALS is often marked by the pronounced formation of cytoplasmic granules, namely TDP43 and FUS granules, we interrogate their formation in S1444P cells. Under normal conditions, no difference was found in cytoplasmic granules (Supplementary Figure 5 A-B). To exacerbate potential effects, we treated cells with two stressors, hydrogen peroxide and sodium arsenite, both of which mimic oxidative stress (Supplementary Figure 5 A-C). Again, no difference was found, suggesting that either granule formation is not a hallmark in the SRRM2 S1444P-induced form of ALS or that HEK 293T cells are an insufficient model to explore this phenomenon.

### A single amino acid exchange in S1444P SRRM2 led to wide-spread transcriptome changes indicative of synapse defects

We therefore focused on cellular pathways that were likely influenced by SRRM2 variants. Due to the importance of nuclear speckles in gene expression and processing, we interrogated the transcriptome of S1444P cells to identify genes and expression patterns that were influenced by this SRRM2 mutation (Figure 4A). SRRM2 wt and S1444P cells clearly segregated in transcriptome analysis (Supplementary Figure 6A) and we could confirm the single nucleotide edit leading to the SRRM2 S1444P (Supplementary Figure 6B). Over 1,300 genes were significantly changed in their expression (Figure 4A, Supplementary Figure 6C, Supplementary Table S3). Of these, comparable numbers were up- and down-regulated, suggesting that these changes are not due to systemic processing defects. We performed gene enrichment analysis to identify affected pathways. Analysis of pathways enriched in downregulated genes with a 1.5-fold cutoff led to several affected pathways (Supplementary Table S4). We restricted our analysis to genes with a 3-fold decrease to increase stringency. We found that genes associated with synaptic transmission and synaptic membrane adhesion were downregulated as well as genes associated with general nervous system processes (Figure 4B). In addition, cytokine response pathways, especially those associated with inflammatory signaling, were also downregulated (Figure 4B). Upregulated genes with a more than 1.5-fold change, on the other hand, were exclusively enriched for positive regulators of synapse formation (Figure 4C). This suggests that even in HEK 293T cells, gene signatures related to synapse formation and function are regulated by SRRM2 and perturbed by the S1444P variant.

**Figure 4.**
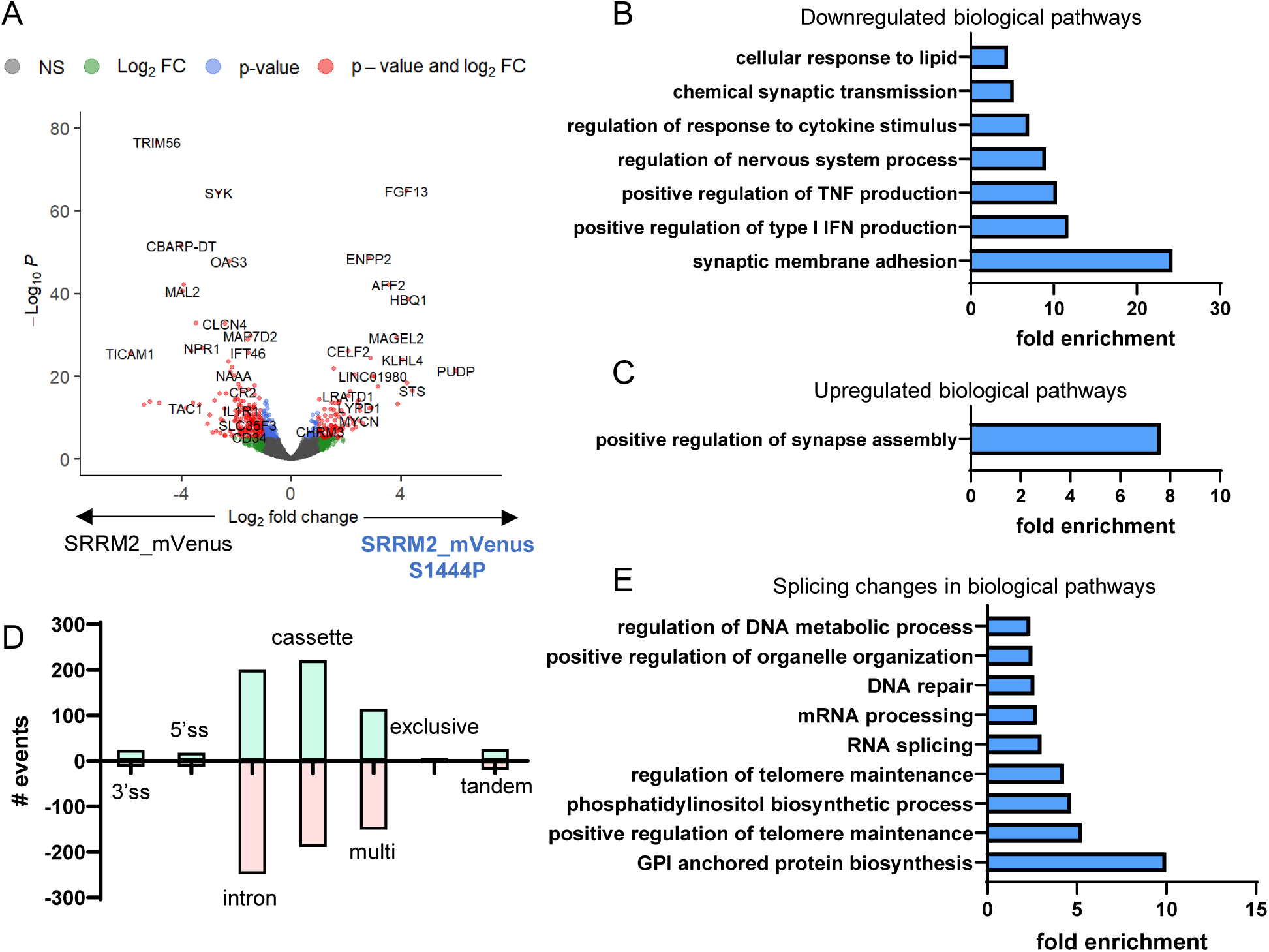
Transcriptome and splicing changes in SRRM2 S1444P cells. (A) Volcano plot of wt and SRRM2 S1444P transcriptomes. The x-axis shows the log fold change (effect size), and the y-axis shows the –log□□ adjusted p-values (significance). (B) Significantly downregulated genes (log□ fold change ≥ 3) were enriched for genes associated with inflammatory cytokines, synapse function, and nervous system processes. (C) Significantly upregulated genes (log□ fold change ≥ 1.5) were enriched for genes associated with the synapse assembly pathway. (D) Splicing differences by events indicated by a dPSI > 0.05 (5%) and a p-value <0.05. (E) Differentially spliced genes were enriched in pathways related to mRNA processing and RNA splicing. 3’ss: 3’ splice site, 5’ ss: 5’ splice site, intron: alternative intron, cassette: cassette exon, multi: multi-exon spanning, exclusive: mutually exclusive exons, tandem: tandem cassette. (A-E) S1444P: amino acid exchange in SRRM2. wt: wildtype. RNA-seq experiments were performed in biological triplicates.

As SRRM2 is a known splicing factor and a constitutive member of the spliceosome, we analyzed mRNA splicing changes. To this end, we used Majiq to quantify splicing events across categories (Figure 4D). We found 422 genes with splicing changes, with alternative intron usage, cassette exons, and multi-exon spanning reads being the most affected event types (Figure 4D, Supplementary Figure 6B, Supplementary Table S5). Generally, the effect size was small, however, differentially spliced genes were enriched for relevant pathways, such as the regulation of DNA repair and telomere maintenance, mRNA splicing and mRNA processing, and, more unexpectedly, membrane protein synthesis (Figure 4E). Overall, this suggests that gene expression changes upon the subtle perturbation of nuclear speckles, could be a molecular driving force of ALS-disease in patients with SRRM2 S1444P.

## Discussion

Here, we identified a new ALS-associated gene, the splicing factor and nuclear speckle scaffold SRRM2. We found that a variant of SRRM2, S1444P, co-segregated with disease phenotypes in an ALS-affected family and the SNV in SRRM2 was predicted to be the most damaging. To assess the molecular underlying, we constructed a cell line to explore the impact of S1444P on SRRM2 localization and interactome, as well as mRNA expression and processing. Interestingly, SRRM2 S1444P lowered the co-localization between SRRM2 and SON, another nuclear speckle factor, and led to the specific loss of interaction with ACIN1. SRRM2 S1444P led to wide-spread gene expression changes, with genes enriched for synapse-associated pathways, suggesting that SRRM2 S1444P could cause alterations in synapse function in affected patients.

Mutations in SRRM2 have been predominantly associated with neurodevelopmental disorders, with symptoms ranging from hypotonia, epilepsy, hyperphagia and obesity, to schizophrenia (28, 42, 43). The majority of these phenotypes were driven by SRRM2 mutations leading to premature termination codons or frameshifts, both of which can abrogate large parts of SRRM2. This suggests that SRRM2 haploinsufficiency could be causal for neurodevelopmental disorders. However, single amino acid exchanges have also been reported (44). Despite its ubiquitous expression, mutations in SRRM2 seem to predominantly affect neural tissues.

In the context of ALS, SRRM2 has been found to be mislocalized in cells with intronic repeat expansions in C9ORF72, where SRRM2 is sequestered in cytoplasmic granules, causing disruption of nuclear speckle properties (6). Further, downregulation of SRRM2 affects TDP-43 mislocalization and loss of function (45). This common association of SRRM2 with known genetic risk factors of ALS, made it likely that genetic mutations in SRRM2 itself could also lead to symptoms within the ALS disease spectrum. Of note, neither TDP43, FUS, nor C9ORF72 were altered in their expression or splicing in our RNA-seq data (Supplementary Table 4).

Synapse loss precedes other symptoms in ALS, which has been termed a synaptopathy, and a distinct loss of synapses can be detected in affected patients (46–48). Interestingly, the exchange of one amino acid in SRRM2 led to a clear gene expression signature, which was enriched in genes associated with the regulation of synapse formation. The striking upregulation of this pathway in HEK cells suggests that it is likely highly conserved. We have previously found that metabolic fluctuations, as found during inflammation, can alter SRRM2 mobility in the nucleus and subsequently gene expression and splicing (19). In this study, inflammatory cytokine pathways, such as IFN and TNF signaling, were modulated in S1444P cells (Figure 4B), confirming a possible connection of SRRM2 with the cellular response to inflammation.

mRNA splicing changes were found in pathways known to buffer and to respond to perturbations in splicing factors, such as feedback to other splicing factors to “buffer” disruptions, as well as to DNA damage response pathways (Figure 4E). mRNA processing and metabolism-associated genes are among the common genetic determinants of ALS. Similar to SRRM2, ACIN1 is also a nuclear protein localized to nuclear speckles (49) and the spliceosome (50), and as such, a likely interaction partner of SRRM2. While ACIN1 (1,341 amino acids) is smaller than the 2,752 amino acid-long SRRM2 (51), ACIN1 is still a large protein. Similar to SRRM2, ACIN1 is mostly disordered with structure prediction suggesting long unfolded stretches interspersed by alpha-helical motifs. As such, it is surprising that the interaction between these two proteins could be, at least partially, disrupted by a single amino acid exchange. The substitution of a serine with proline removes simultaneously a serine phosphorylation site (52) and could cause bends and kinks in secondary structures (53). It is of note that disordered regions can still undergo precise interactions, often driven by charge distributions, adaptation of an ensemble of conformations, fuzzy binding through distinct individual binding states, or folding upon binding (54). SRRM2 phosphorylation determines the physical properties of nuclear speckles, and expression of SRRM2 variants that either cannot be phosphorylated or mimic constitutive phosphorylation, can change mRNA splicing and DNA repair, respectively (55). While we did not find SRRM2 S1444P phosphorylation in our dataset, which is likely due to the lack of enrichment of phosphopeptides (Supplementary Table S6), its post-translational modification was confirmed in other studies (52, 55).

Due to its unusual amino acid sequence, SRRM2 has an interesting dual role as a nuclear speckle scaffold and a constitutive part of the active spliceosome. Due to its long, disordered C-terminus, SRRM2 offers both ample sequence space for interactions as well as regulation through phosphorylation and other post-translational modifications. As such, SRRM2 could provide an interaction interface for other splicing factors to position themselves in the proximity of active spliceosomes nearby and in nuclear speckles. Other reported amino acid exchanges could lead to a similarly precise change in the SRRM2 interactome, while consequences of SRRM2 termination and frameshifts might lead to a more general disruption of nuclear speckle architecture.

## Material and Methods

### Exome capture and next-generation sequencing

Exome enrichment of 3 mg genomic DNA was performed by using the Agilent SureSelect Human All Exon Kit capture library G3370 according to the manufacturer’s protocol (SureSelect Human All Exon Illumina Paired-End protocol version 1.0.1). The capture library, containing regions totaling approximately 50 megabases, is designed to target all human exons in the NCBI consensus coding sequence (CCDS) database. Captured DNAs were sequenced on an Illumina GAIIx sequencer (Illumina) with 100-base-pair paired-end reads. Multiple lanes of sequencing data were collected for each sample to generate sufficient coverage. Image analyses and base calling were performed by using the Illumina Genome Analyzer Pipeline software (GAPipeline version 1.5 or higher) with default parameters. Reads were aligned to a human reference sequence (UCSC assembly hg19, NCBI build 37) and genotypes were called at all positions at which there were high quality sequence bases (Phred-like score Q25 or greater) at minimum coverage of 8, using CLC Genomics Workbench v4.5.1 (CLC Bio). For each sample, at least 180 million reads from each sample were uniquely mapped to the targeted human exon regions to give a mean depth of coverage of 240. Under such coverage, approximately 95% of targeted regions were covered by 10 reads or more, and more than 86% were covered by more than 40 reads. To identify the pathogenic mutations, we performed a series of filtering steps (Supplementary Fig. 1). We first discarded the variants that did not change the amino-acid sequence. We then generated a list of variants that are present in affected individuals but absent in nonaffected individuals. Variants previously reported in the Single Nucleotide Polymorphism Database build 132 (dbSNP132) hosted by the National Center for Biotechnology Information were excluded. The variants were further filtered by using the in-house database (a collection of variants from whole-exome sequencing of more than 625 individuals) with a similar methodology. Finally, Sanger sequencing was performed on DNA of three additional family members (two affected and one non-affected) to exclude nonpathogenic mutations.

### Cell culture

HEK 293T cells were obtained from ATCC and confirmed by STR analysis. Cells were cultured at a density of 80-90% confluency and passaged twice per week. If not stated otherwise, cells were maintained in high glucose Dulbecco’s modified Eagle’s medium (DMEM; Sigma-Aldrich, D5796), with 10% Fetal Bovine Serum (FBS; Corning, 35-007-CV), and 1% Penicillin/Streptomycin (Gibco, 15140-122). Cells were tested monthly for mycoplasma contamination and were consistently found negative. Experiments were performed within passage 20.

### Prime editing and single cell sorting

The PEA1-mCherry plasmids were adapted from PEA1-GFP plasmids through PCR and cloned with guide RNA (gRNA), repair template, and second nick gRNA designed using PETAL. Repair template sequence was then optimized with 3’ motifs using PegLit. The plasmid was co-transfected into HEK 293T cells with pCMV-PE6c using CalFectin (SignaGen). Positive cells were selected based on mCherry expression by the TEMERTY Faculty of Medicine flow cytometry facility using BD FACSAria™ **I**u Cell Sorter. Cells were sorted as single cells into each well of a 96-well plate supplemented with high glucose DMEM with 20% FBS and 1% Penicillin/Streptomycin. The cells were then expanded to a 24-well plate for follow-up screening by Sanger sequencing. PEA1-GFP was a gift from Paul Thomas (Addgene plasmid #171993) (36), pCMV-PE6c was a gift from David Liu (Addgene plasmid #207853) (37).

### Sanger sequencing

DNA was isolated from the cells at 80-90% confluency using DNA extract solution containing proteinase K (QuickExtract DNA Extraction Solution, Lucigen). The primers were designed to amplify 596 bp surrounding the T. 4089C site in a three-step PCR using Phusion polymerase (NEB, M0530S). The molecular weight of the amplified product was confirmed with 2% agarose gel electrophoresis. The product was then purified with the Monarch Spin PCR & DNA Cleanup kit (NEB, #T1030L). 100 ng of DNA were sent for Sanger sequencing with a custom primer at The Centre for Applied Genomics Facilities in the SickKids hospital, Toronto.

### RNA isolation

Cells were seeded two days prior to the extraction at a density of 0.5 × 10^6^ cells/well in a 6-well plate. After washing with PBS (Sigma Aldrich, D8537) for two times, RNA was extracted and purified with the PureLink^TM^ RNA mini kit (Invitrogen, 12183018A) following the manufacturer’s instructions. RNA integrity was confirmed by 28S/18S rRNA band ratio in 1% agarose gel electrophoresis and concentration was measured with NanoDrop.

### RT-qPCR

The cDNA was synthesized from 1 µg of Total RNA using the RevertAid First Strand cDNA Synthesis Kit (Thermo Scientific, K1621) with Oligo-dT according to the manufacturer’s instructions. The real-time quantitative PCR (qPCR) was performed using the LightCycler® 480 System (Roche). Amplification was performed in a 25 µL reaction containing 12.5 µL 2x PowerUp SYBR Green Master (Applied Biosystems, A25742), 500 nM forward and reverse primers, 20 ng of cDNA (added up to volume with Nuclease-Free Water). Cycling conditions were 95°C for 10 mins, followed by 50 cycles of 95°C for 15 s, 60°C for 30 s and 72°C for 30 s. Each sample was run in technical triplicates and with biological replicates n ≥ 3. Expression was normalized to GAPDH, and the fold change was calculated with 2^ΔΔCt method. All the qPCR primers used are in the Supplementary Table S8.

### Immunofluorescence

293T SRRM2-mVenus and S1444P mutant cells were grown on poly-D-lysine (Sigma-Aldrich, P7280-5MG) coated coverslips (Electron Microscopy Science, #72230-01, 12mm). The samples were rinsed with PBS and fixed with 4% PFA (BioShop, PAR070) for 15 mins at RT. The fixed cells were then permeabilized with 0.25% Triton X-100/PBS for 5 mins RT, and following by blocking with 1% BSA (BioShop, 9048-46-8) and 2% goat serum (Corning) in PBS for 1 hour at RT. Primary antibodies targeting SON or ACIN1 were diluted in blocking buffer and incubated with the cells overnight at 4°C. Next day, cells were washed in blocking buffer 3 times for 5 mins and incubated with DyLight 650 secondary antibody (1:200) for 90 mins at RT prevented from light. To counterstain DNA, cells were incubated with DAPI (Roche, 10236276001) for 15 min at RT, and then washed 3 times with PBS. Coverslips are briefly rinsed with distilled water and mounted on glass slides using ProLong Gold Antifade Reagent (Cell Signaling Technologies, #9071S) overnight at RT. Samples were sealed with nail polish and stored at 4°C in dark chamber before imaging. All antibodies are listed in Supplementary Table S7.

### Confocal Microscopy

Images were acquired on a Leica TCS SP8 confocal microscope (Leica) with 63×/1.40 Oil HC PL APO CS2 objective and LAS X software. Coverslips were mounted with Type F Immersion Liquid (Leica). Images were taken at 1024 x 1024 with identical acquisition settings across samples, analyzed with ImageJ/Fiji and exported as TIFF files.

Colocalization analysis was performed using Imaris (Oxford Instruments) on individual cell z-stacks, and Manders coefficients were calculated for each cell.

### Cell proliferation assay

293T SRRM2-mVenus and S1444P mutant cells were seeded into a 96-well tissue culture plate (Sarstedt) with a cell density of 5 x 10^3^ cells/well. Proliferation was assessed with AlamarBlue reagent (Invitrogen, DAL1025) added at 1:10 in DMEM, and plates were incubated for 2 hours at 37°C. Fluorescence was recorded on a microplate reader (Synergy H1, BioTek) with λ_ex_ = 560 nm, λ_em_ = 585 nm and gain held constant across plates. One plate was measured every 24 hours for a total of 5 days (0-96 hours, in 24 hours intervals). Background signals from DMEM + AlamarBlue were measured and subtracted, and the final values were normalized to Day 0 (t=0). All samples were run in quadruplets and in biological replicates n ≥ 3.

### Induction of cell stress

293T SRRM2-mVenus and S1444P mutant cells were grown on poly-D-lysine coated coverslips. Cells were treated with hydrogen peroxide (200 µM or 1 mM) for 15 mins to 1 hour, or sodium arsenite (0.1 mM or 0.5 mM) for 1 hour. Cells were then rinsed with PBS and fixed as indicated above. Primary antibody targeting TDP43, FUS or G3BP1 were diluted in blocking buffer and incubated with cells overnight at 4°C. Next day, cells were rinsed and incubated with AlexaFluor 488 secondary antibody (1:100 - 1:200) for 90 mins at RT prevented from light. To counterstain F-actin, cells were incubated with AlexaFluor 647 Phalloidin (Cell Signaling, 8940S) for 20 minutes at RT, followed by DAPI (Roche, 10236276001) for 15 min at RT. All antibodies are listed in Supplementary Table S7.

### Flow cytometry

293T SRRM2-mVenus and S1444P mutant cells were collected at 80-90% confluency and filtered through 70 µM cell strainers. Cells were analyzed on the Cytek Aurora (3 lasers, violet (405 nm), blue (488 nm), and red (640 nm)). A minimum of 20,000 events were collected per sample. Debris and doublets were excluded by forward and side scatter gating. Data was analyzed using FlowJo software (BD Biosciences). Fluorescent intensities were quantified as mean fluorescence intensity (MFI)

### Protein isolation

293T SRRM2-mVenus and S1444P mutant cells were collected at 80-90% confluency and lysed with 1x RIPA lysis buffer (0.05 M Tris/HCl, pH = 7.4, 0.15 M NaCl, 0.25% deoxycholate, 1% NP-40, 1 mM EDTA) with cOmplete Mini Protease Inhibitor Cocktail (Roche, 11836170001). Lysed cells were spun at 14,000 x g for 10 mins and the supernatant was collected. Samples were prepared with 5x SDS loading dye and boiled at 95°C for 10 mins. If not using immediately, samples were stored at -20°C.

### GFP nanobody purification

Plasmids encoding GFP-nanobody were a gift from Brett Collins (Addgene #49172) (38). The GFP-nanobody sequence was cloned into pET28b_SUMO and expressed as a SUMO-nanobody fusion protein overnight at room temperature in LB medium in BL21 *E. coli* Cells were lysed using a microfluidizer in 40 mM Hepes, pH=8.0, 500 mM NaCl, 50 mM imidazole) and His-tagged SUMO-nanobody was bound to Ni-NTA agarose (Qiagen) through gravity flow. Beads were washed with lysis buffer and protein was eluted by the stepwise addition of increasing concentrations of lysis buffer (40 mM Hepes, pH=8.0, 500 mM NaCl, 500 mM imidazole end concentration). Fractions containing the nanobody were identified with SDS PAGE, concentrated, and loaded on a Superdex 200 size exclusion column (Cytiva). Following confirmation of purity by SDS PAGE (∼90%), the resulting nanobody was concentrated to 2 mg/ml and frozen at - 80°C until use. Magnetic CNBr Activated High Flow Beads (ProteinArk) were coupled to the purified nanobody according to the manufacturer’s instructions. Beads were stored in PBS at 4°C and used within 1 month of coupling.

### Co-Immunoprecipitation

293T SRRM2-mVenus and S1444P mutant cells were seeded onto 15 cm dishes at a density of 5 × 10^6^ cells per dish and incubated at 37°C for 48 hours. Cells were harvested in 1 mL of mild lysis buffer (Tris-buffered Saline with 1% NP-40 and 1 mM MgCl_2_) supplemented with Pierce Phosphatase Inhibitor (Thermo Scientific, A32957) and cOmplete Mini Protease inhibitor cocktail (Roche, 1183670001). Lysates were incubated at 4°C for 30 mins, followed by centrifugation at 14,000 × g for 20 mins. Supernatant was collected and incubated with 90 µL of Magnetic-Coupled GFP nanobody overnight at 4°C. Next day, beads were collected with magnets and washed 3 times with wash buffer (50 mM Tris-HCl pH 7.4,150 mM NaCl, 0.05% NP-40). For western blot, bound protein was then eluted with 2x SDS loading dye and boiled at 95°C for 10 mins. For Mass spectrometry, beads were stored at -80°C in 90 µL PBS prior to mass spectrometry.

### Interactome sample preparation

Beads were incubated with 25 µL Elution buffer I (5 ng/µL trypsin, 2 M urea, 50 mM Tris-HCl pH 7.5, 1 mM DTT) at RT for 30 mins, followed by adding 100 µL Elution buffer II (2 M urea, 50 mM Tris-HCl pH 7.5, 5 mM chloroacetamide) and incubating at RT for 16 hours with gentle agitation. Next day, samples were quenched with 1% formic acid and the supernatant was desalted using C18 StageTips (Pierce) according to the manufacturer’s protocol. After washing with 0.5% formic acid, peptides were eluted in 30 µL of Elution Buffer III (80% acetonitrile/0.5% formic acid), speedvac concentrated (Thermo Fisher Scientific) and stored dry at -80°C until the experiment. Prior to analysis, peptides were resuspended in 40 µL of 1% formic acid and analyzed using an EASY-nLC 1200 UHPLC system combined with a Q-Exactive HF-X orbitrap mass spectrometer via an in-line nanoLC-electrospray ion source (Thermo Fisher Scientific). Solubilized peptides were loaded onto a house-made fused-capillary silica precolumn (100 μm I.D.) packed with 2 cm of 5 μm Luna C18 100 Å reverse phase particles (Phenomenex) and separated on a 14.1 cm (100 μm I.D) silica pulled emitter packed with 1.9 μm Luna C18 100 Å reverse phase particles (Phenomenex); capillary was sourced from Polymicron Technologies. Peptides were eluted over 120 mins with mobile phase A (0.1% FA) and mobile phase B (80/20/0.1 ACN/water/FA) at a constant flow rate of 300 nL/min with a linear increase from 2% to 5% B over 1 min, an increase to 26% B over 70 mins, an increase to 60% B over 20 mins, then by an increase to 100% B over 14 mins, and a final plateau at 100% B for 15 mins.

### Mass spectrometry data acquisition and analysis

Mass spectrometry was performed, as described by Hall, Yeung, and Peng (2020) (56). In brief, a data-dependent top 20 method was used. Full MS scans were acquired from *m/z* 350–1400 at a resolution of 60,000 at 200 *m/z* with a target automated gain control (AGC) of 3 × 10^6^ charges. For higher-energy collisional-dissociation MS/MS scans, the normalized collision energy was set to 28 and a resolution of 15,000 at 200 *m/z*. Precursor ions were isolated in a 1.4 *m/z* window and accumulated for a maximum of 20 ms or until the AGC target of 1 × 10^5^ ions was reached. Precursors with unassigned charge states, a charge of 1+, or a charge of 7+ and higher were excluded from sequencing. Previously targeted precursors were dynamically excluded from re-sequencing for 20 s.

Mass spectrometry data were processed with MaxQuant 2.6.4.0 and Perseus 2.1.2 (57, 58) as described previously (59) to identify IleRS interaction partners. In brief, raw files were searched against Homo sapiens reference proteome UP000005640_9606 (60) (Uniprot) with the Andromeda search engine integrated into MaxQuant: M oxidation, N-terminal actylation as variable, C carbamidomethyl as fixed modification. 2 missed cleavages maximum. First search peptide tolerance 20 ppm, main search peptide tolerance 4.5 ppm. Protein False Discovery Rate set to a maximum of 0.01 (1%). Minimum peptides 1. Default settings were used for label free quantification (LFQ, minimum ratio count 2). The resulting protein groups file was loaded into Perseus and filtered for ‘reverse’, ‘potential contaminants’ and ‘only identified by site’. The log2 values of LFQ intensities were calculated, and all proteins with less than two valid values/group were discarded. Missing values were replaced from a normal distribution (width 0.3 and downshift 1.8), and unpaired, two-tailed Students t-test was used to calculate significance and difference.

### RNA-seq library preparation

Total RNA samples with RNA integrity number scores > 8, as assessed by Bioanalyzer (Agilent), were used for library preparation. Libraries were generated using NEBNext Ultra II RNA Library Prep Kit for Illumina (NEB) in combination with NEBNext Multiplex Oligos for Illumina (Dual Index Primer Set I) (NEB), following the manufacturer’s protocol. Briefly, poly(A) mRNA was isolated from 1 µg of total RNA input using oligo dT magnetic beads, fragmented, and reverse-transcribed to synthesize first-strand cDNA. Second-strand cDNA synthesis was performed to generate double-stranded cDNA. The resulting cDNA product was subjected to end repair/dA-tailing and adaptor ligation. Libraries were PCR-amplified with unique dual index primers for 7 cycles and purified with NEBNext Sample Purification Beads. Libraries were assessed for quality and concentration, and sequencing was performed by the Donnelly Sequencing Centre at the University of Toronto.

### RNA-seq Analysis

>50 million read pairs were sequenced as 150 nt paired end reads for each of three biological replicates by the Donnelly Sequencing Core using a Novaseq6000 SP 300C. Reads were quality controlled and adapter trimmed using Trimmomatic 0.6.4 (61) and mapped to Homo_sapiens.GRCh38.114 (62) using STAR aligner 2.7.11b (63). Counts were obtained using RSubread’s FeatureCount function (Version 2.12.3) (64) and differentially expressed genes were identified with DESeq2 1.38.3 (65). Splicing changes were interrogated with Majiq and Voila V3 (66, 67).

### Statistics

Significantly enriched interactors were identified using unpaired, two-tailed Students t-test in Perseus. Differentially expressed genes were determined using DESeq2, padj < 0.05. Differentially spliced genes were identified using Majiq/Voila, with sites filtered for a change in percent spliced in (PSI)>5% and a pvalue <0.05.

## Data Availability

Antibodies and oligonucleotides used throughout the study are listed in Supplementary Table 7 and 8.

Material is available upon request to.

## Acknowledgements

We acknowledge the Centre for Applied Genomics at SickKids Hospital for Sanger sequencing. We thank the Temerty Faculty of Medicine Flow Cytometry Facility for cell sorting, the Cell and Systems Biology Imaging Facility, and the Donnelly Sequencing Centre, all at the University of Toronto. We also greatly appreciate access to flow cytometry through Prof. Landon Edgar’s group at the University of Toronto, Faculty of Medicine, Department of Pharmacology and Toxicology. HC and CL thank Dr. Colin Campbell at the University of Edinburgh for supporting CL’s thesis.

## Data availability

Material is available upon request to.

## Author contribution

Conceptualization: QS, JPT, HC. Data curation: QS, YDW, HC. Formal analysis: QS, CL, YDW, JPT, HC. Funding acquisition: JPT, HC. Investigation: QS, CL, JMM, YDW, NE, SV, BDF, HC. Methodology: QS, CL, JMM, YDW, NE, SV, BDF. Resources: JPT, HC. Supervision: HP, HJK, JPT, HC. Visualization: QS, HJK, HC. Writing – original draft: QS, HJK, HC. Writing – review & editing: QS, JMM, BDF, HP, JPT, HC.

## Supplementary Table legends

Table S1: SRRM2 wt interactome.

Table S2: SRRM2 S1444P interactome.

Table S3: Differentially expressed genes S1444P vs wt.

Table S4: GO analysis of pathways enriched with differentially expressed genes.

Table S5: Splicing comparison S1444P vs wt.

Table S6: Phosphopeptides identified in SRRM2 wt and S1444P mass spectrometry.

Table S7: Antibodies used in this study.

Table S8: Oligonucleotides used in this study.

---

**Figure S1.**
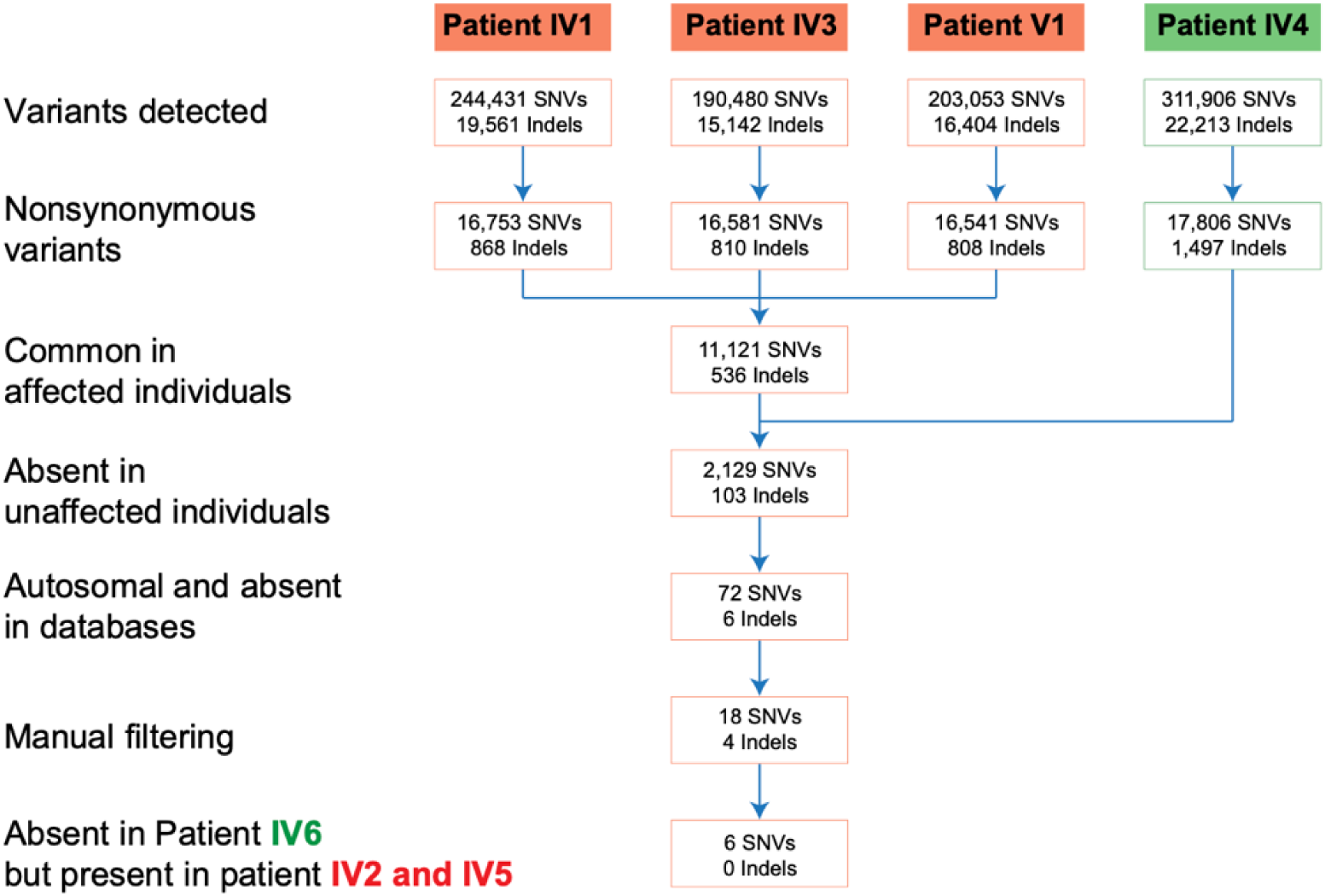
Tissue expression of segregating genes and interactome overlap of ALS-associated proteins. The filtering paradigm used to detect novel sequence variants that co-segregated with disease is shown. Affected patients are represented by red and unaffected patients by green.

**Figure S2.**
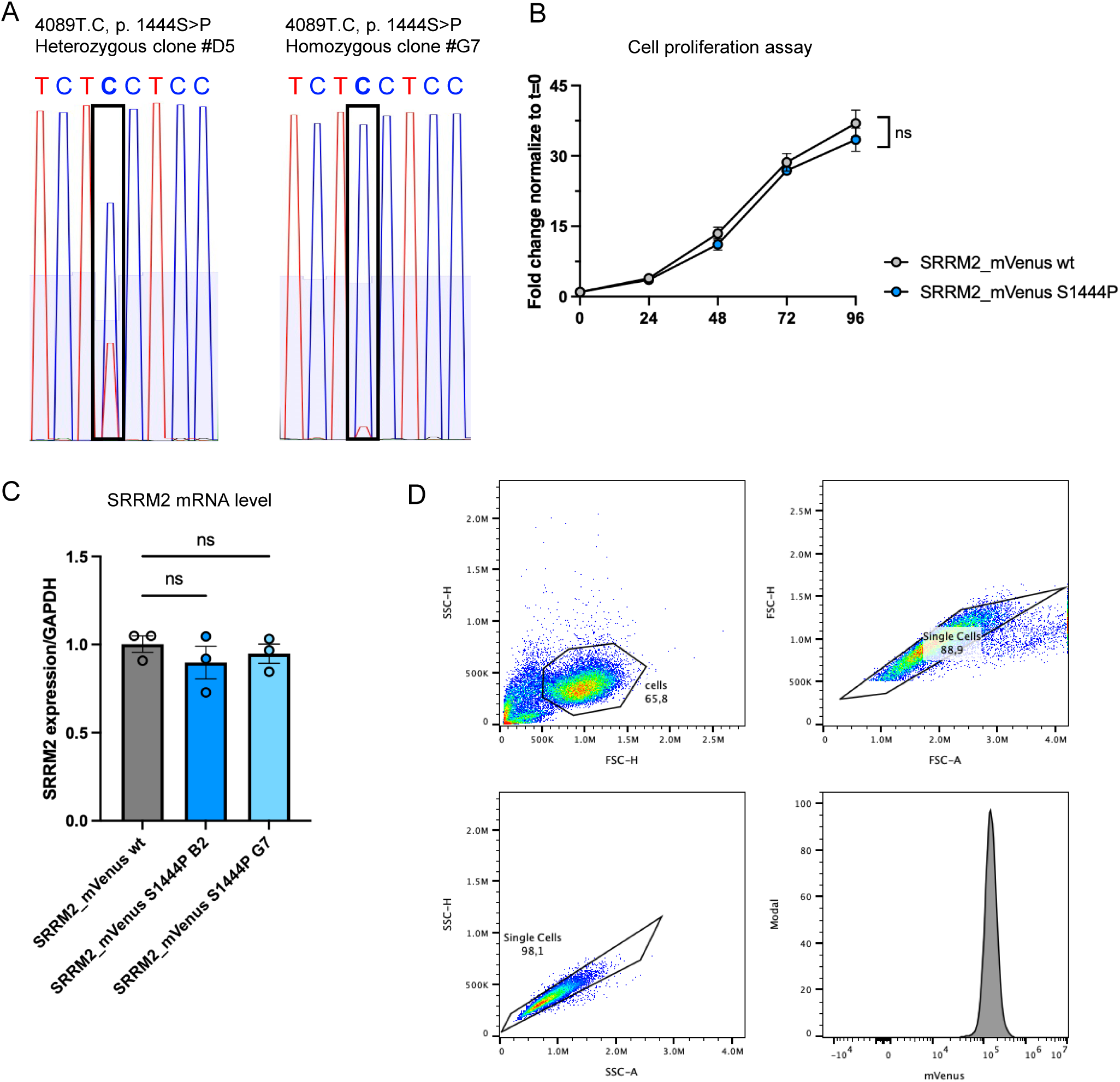
Characterization of SRRM2 S1444P model cell lines. (A) Nanopore sequencing to confirm heterozygous and homozygous SRRM2 4089T.C (S1444P) cell lines. (B) Cell proliferation, measured by Alamar blue fluorescence, showed no difference between wt and S1444P cells. A representative of at least 3 replicates is shown. (C) Additional qPCR to assess SRRM2 mRNA levels following gene editing. (D) Representative gating scheme used for data analysis. Cells were first gated on FSC-H versus SSC-H to exclude debris, followed by doublet exclusion using FSC-H versus FSC-A and SSC-H versus SSC-A. The MFI of mVenus was then quantified in the appropriate detection channel. FSC: forward scatter, SSC: side scatter, H: Height, A: Area, MFI: mean fluorescent intensity.

**Figure S3.**
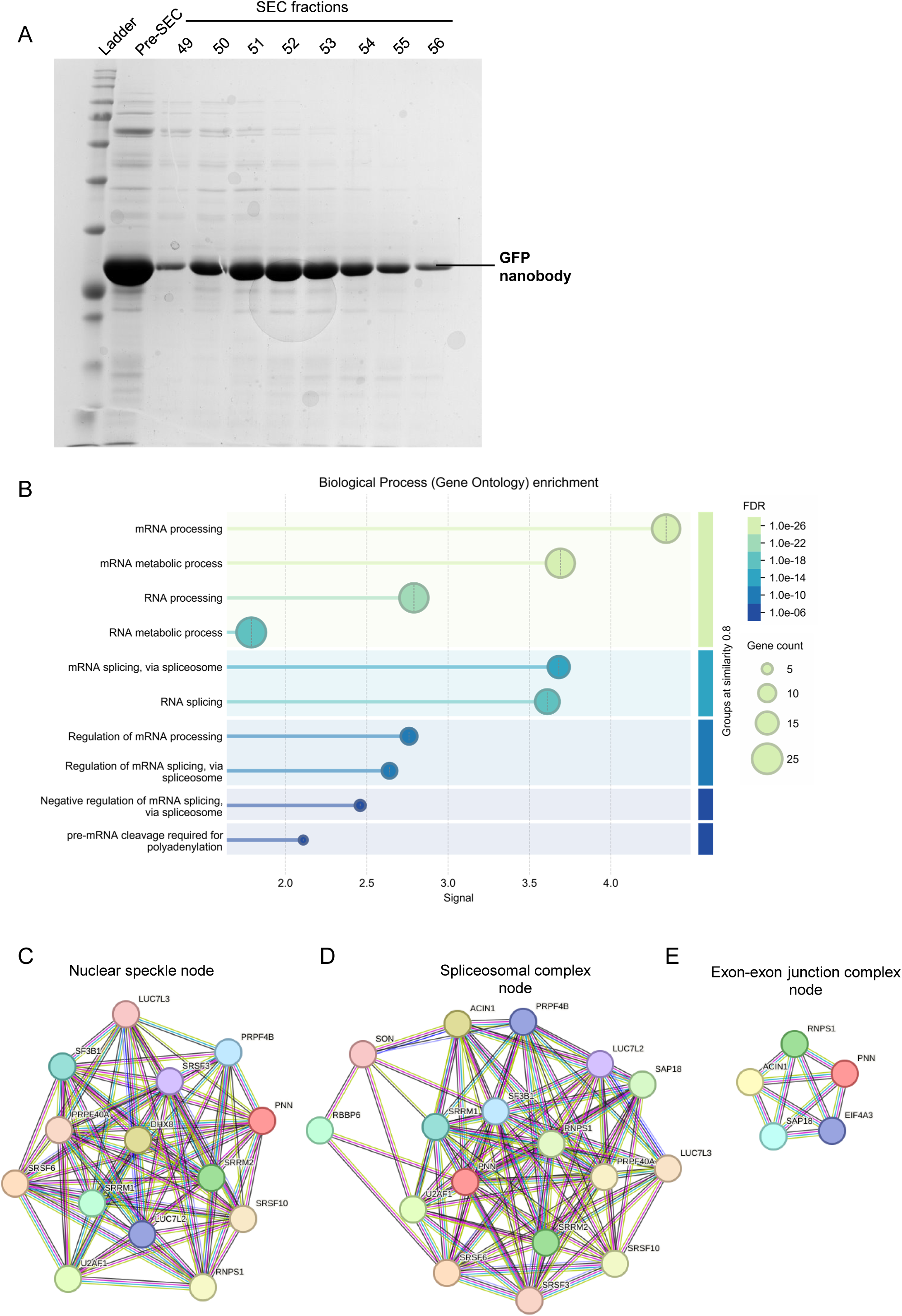
SRRM2 wt and S1444P interactome analysis. (A) SDS-PAGE confirming purification of the GFP-nanobody, which was subsequently used for enrichment of SRRM2_mVenus. SEC: size exclusion chromatography. (B) Enrichment analysis of the SRRM2 wt proteome confirming that the majority of SRRM2 interactors were nuclear speckle proteins and/or associated with mRNA processing. (C-E). Nodes displaying proteins associated with different complexes in the SRRM2 interactome.

**Figure S4.**
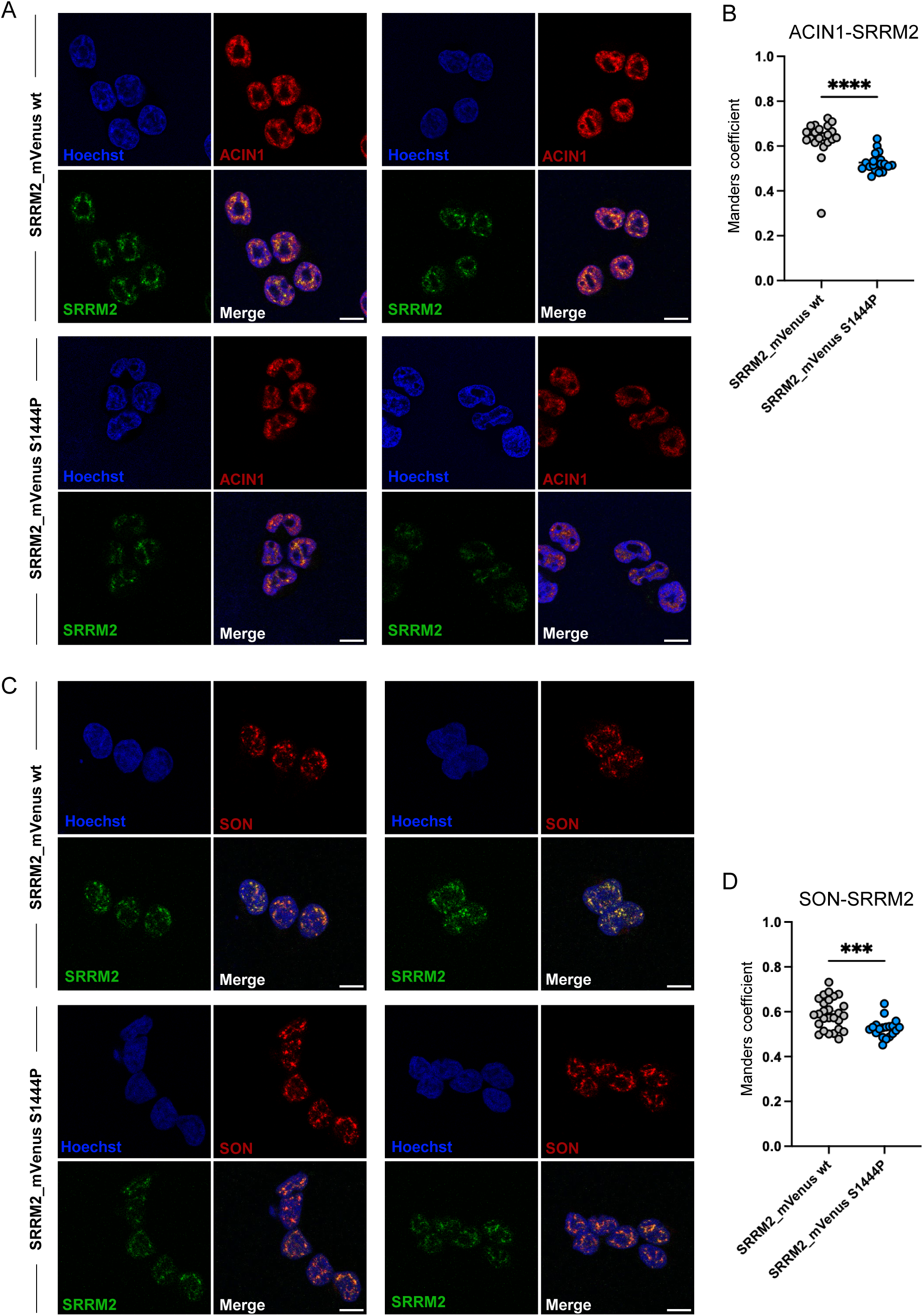
Investigation of stress granule formation in SRRM2 S1444P cells. (A) Additional representative images used to assess SRRM2 and ACIN1 colocalization. SRRM2 is imaged through fusion to the fluorescent protein mVenus, ACIN1 is visualized through immunofluorescence. (B) Manders coefficient of ACIN1 with SRRM2. n=21, 20. (C) Additional representative images of SRRM2 and SON colocalization. SRRM2 is imaged through fusion to the fluorescent protein mVenus, SON is visualized through immunofluorescence. (D) Manders coefficient of SON with SRRM2. n=27, 19.

**Figure S5.**
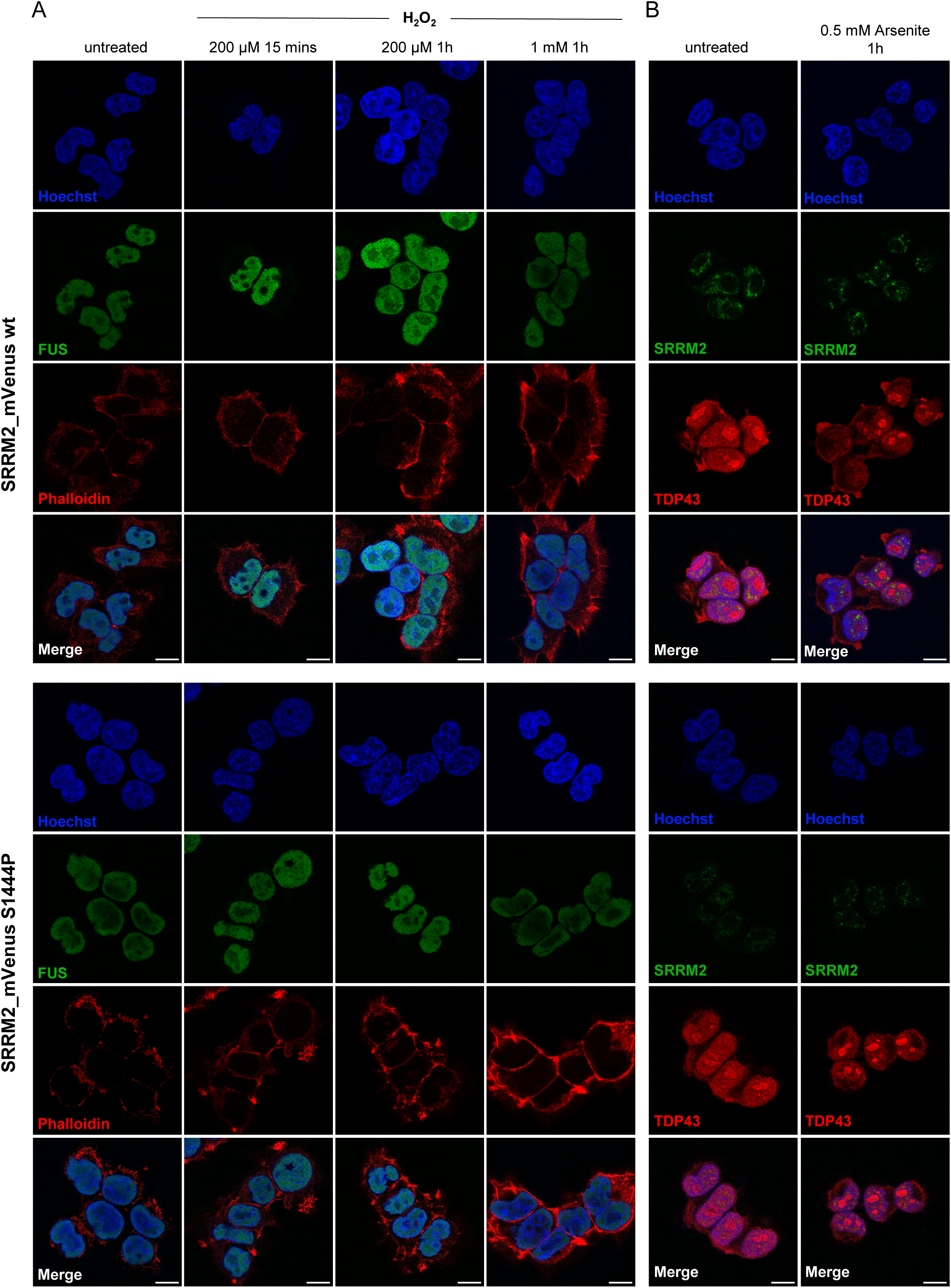
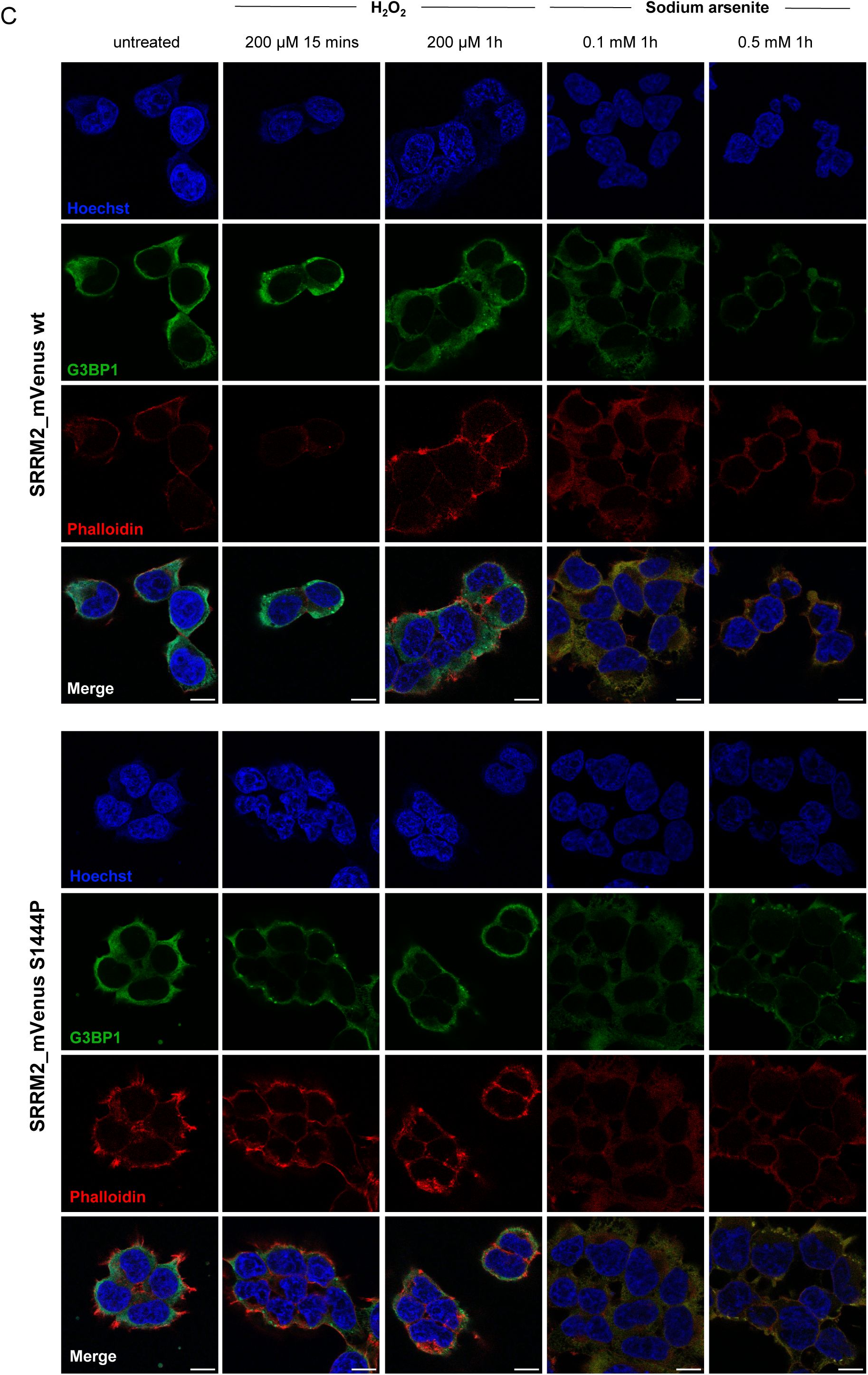
Analysis of stress granule formation in S1444P cells under oxidative stress. (A) FUS immunofluorescence to monitor FUS nuclear export and cytoplasmic aggregation in S1444P cells. Cytoplasmic localization was not observed in wt or S1444P cells. (B) TDP43 immunofluorescence to assess TDP43 mislocalization in S1444P cells. TDP43 localization was comparable between wt and S1444P cells. (C) G3BP1 immunofluorescence indicating stress granule formation in wt and S1444P cells. G3BP1 granule formation was unchanged between wt and S1444P cells but responsive to varying concentrations of stressors. (A-B) A representative out of at least 2 replicates is shown. (C) A representative out of at least 3 replicates is shown. Hoechst labels DNA in nuclei, phalloidin binds F-actin and is used here to indicate cell boundaries.

**Figure S6.**
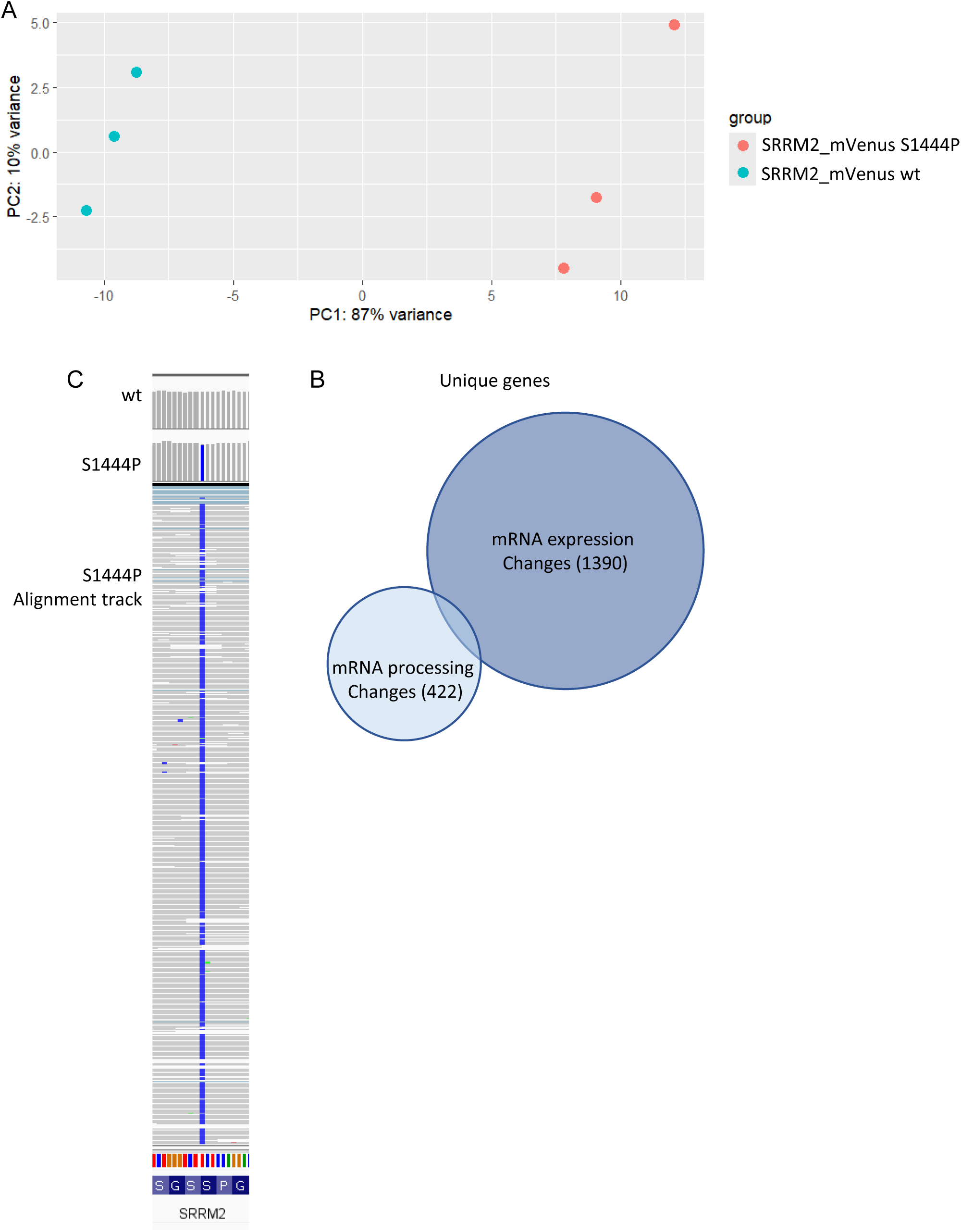
Characterization of SRRM2 S1444P model cell lines. (A) PCA plot showing distinct differences between wt and S1444P transcriptomes. (B) Comparison between differentially expressed and differentially spliced genes in SRRM2 S1444P vs wt cells. (A-B) RNA-seq was performed in biological triplicates. Differentially expressed genes were defined by a padj < 0.05 and a > 2.2-fold change. Differentially spliced genes were defined by a p-value < 0.05 and a dPSI > 0.05.

